# Decision confidence as a mapping of bayesian posterior belief

**DOI:** 10.1101/124891

**Authors:** Luciano Paz, Alejo Salles, Mariano Sigman

## Abstract

We study the confidence response distributions for several two alternative forced choice tasks with different structure, and assess whether their behavioral responses are accurately accounted for as a mapping from bayesian inferred probability of having made a correct choice. We propose an extension to an existing bayesian decision making model that allows us to quantitatively compare the relative quality of different function mappings from bayesian belief onto responded confidence. We find that a simple linear rescaling from bayesian belief best fits the observed response distributions. Furthermore, the parameter values allow us to study how task structure affects differently the decision policy and confidence mapping, highlighting a dissociable effect between confidence and perceptual performance.

## 1 Introduction

Perceptual decision-making based on noisy and variable perceptual stimuli is usually thought to be carried out as the on-line integration of perceptual evidence until an admissibility threshold is reached (Bogacz et al., 2006, Forstmann et al., 2016, Ratcliff and McKoon, 2008, Smith, 2000, Usher and McClelland, 2001). Surprisingly, these models that are natural constructs for time varying stimuli can also account for most behavioral observations derived from decision making based on discrete, static stimuli (Ratcliff and Smith, 2004, 2010, Smith et al., 2014), such as a number comparison task (Sigman and Dehaene, 2005). Even though in static tasks the nature of the sequential accumulation is not clear, these models are most often used to account dynamic and static tasks, although the parameter values are slightly different across task modalities (Ratcliff and Smith, 2010, Smith et al., 2014).

The most typical observations used to constrain and fit models of decision making are response time (RT) distributions and task performance. Another relevant observable that has recently received much attention is confidence, i.e. the subject’s internal measure of how sure he/she is that the reported decision was correct (Kepecs et al., 2008, Yeung and Summerfield, 2012, Zylberberg et al., 2012). There is still not an established consensus on how confidence is encoded and read-out in sensory integration (Kiani and Shadlen, 2009, King and Dehaene, 2014, Ma et al., 2006, Paz et al., 2016, Pleskac and Busemeyer, 2010, Pouget et al., 2016, Vickers et al., 1985, Zylberberg et al., 2012) and, more importantly, how a common encoding could be implemented for both static and dynamic stimuli tasks. For static tasks most confidence models rely on signal detection theory measures (Fleming and Lau, 2014, Grimaldi et al., 2015, Maniscalco and Lau, 2012) whilst for dynamic tasks confidence is modeled using several statistics on the accumulators of sensory evidence (Moreno-Bote, 2010, Vickers et al., 1985, Zylberberg et al., 2012), the most widely used being the logit function of the probability of having decided correctly (Kiani and Shadlen, 2009, Kiani et al., 2014, Zylberberg et al., 2014). A related current discussion is whether the same circuits that encode information for choice also, and at the same time, encode information that is read-out to convey confidence judgments. The evidence in favor of circuit sharing mostly comes from experiments in macaques that show that the same neurons that are thought to encode choice formation also, and at the same time, affect confidence (Kiani and Shadlen, 2009, Kiani et al., 2014). However, if confidence is merely encoded in the same circuitry as choice it is unclear how there are several experiments that show a clear dissociation between decision accuracy and subjective perception (Graziano and Sigman, 2009, Graziano et al., 2015, Zylberberg et al., 2014).

Our objective is two-fold, first, to asses theoretically whether a specific readout of an integration model can explain choice and confidence in static and dynamic tasks. Second, specifically, to ask what mapping (or readout) of the integration process accounts for the observed confidence reports.

We use a bayesian decision model (Drugowitsch et al., 2012, 2014a,b, 2015) that is able to account for behavioral data from both static and dynamic tasks. Furthermore, we propose an extension that aims to provide a normative way to model subjects’ confidence reports using the bayesian inferred probability that the selected action was correct (Drugowitsch et al., 2015). A particular strength of the model is that it can explain the covariance between several observables of decision making such as response time (RT), performance and confidence distribution statistics, and it can relate them to parameters which are hidden from direct observation such as the internal sensory noise or the subject’s urgency. Furthermore, the model parameter values allow us to analyze tasks and sessions similarities across subjects, and we find distinct parameter hierarchies for those directly associated to decision commitment and the parameters linked with the scaling of confidence reports.

With our modeling approach we can ask:

1. Can the extended model account for confidence distributions and their covariance with RTs and performance?
2. How are the model beliefs better converted into confidence judgments?

Furthermore, estimating the model parameters for different individuals, different sessions and different tasks, we can also ask:

3. Is there a correlation structure between the model parameters or are they completely independent from each other?
4. How do the model parameters vary across tasks and sessions, and what do they tell us about the subject’s decision-making policy in different environments?

### 1.1 Background

Decision making for time varying stimuli must in some way assess the probability that each of the available actions is correct based on a stream of evidence samples {*δx*_0_, *δx*_1_, … *δx_T_*}, where we note *δx*_*t*_ as the evidence the observed in the interval [*t*,*t* + *δt*). Then, an admissibility level must be set, which indicates when it becomes better to commit to a decision based on a behavioral goal.

The model we use aims to study two alternative forced choice (2AFC) tasks in which each of the available actions can be mapped to an hypothesis that underlies the generation of the observed evidence, and thus it becomes necessary to infer the probability of each hypothesis given the observed stream (we note the belief that a hypothesis *H_i_* is correct as *g*_*i*_). The model achieves this using Bayes rule 
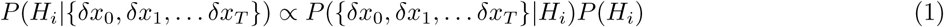
where *P*(*H*_*i*_) is hypothesis *H*_*i*_’s prior probability, and *P*({*δx*_0_, *δx*_1_, … *δx_T_*} | *H_i_*) is the likelihood of the observed evidence given H_i_.

The model computes the admissibility level to commit to a decision by attempting to maximize its perceived reward rate. This implicitly assumes that subjects perceive some form of reward after successfully completing a trial, that the passage of time affects the total perceived reward, and thus they attempt to balance their performance with the time they take to respond. Drugowitsch et al. (2012) showed for 2AFC how to compute the time varying bounds for the belief, *g*, of each alternative that yielded maximal reward rate even in presence of a cost for the passage of time *c*(*t*). It was also shown that the belief bounds could be mapped to the accumulated evidence *x*(*t*) = ∫ *δx_t_dt*, which for certain generative models behaved as a diffusing particle with drift. Using the computed bounds, the generative model and adding a random non-decision time after reaching the bounds, it is possible to compute the model’s predicted RT distribution and performance.

We extend Drugowitsch et al. (2012) optimal decision making model to use the belief that the selected hypothesis is correct at decision time to produce confidence reports. We assume that confidence is reported on a continuous interval [0, 1] where 0 and 1 are the lowest and highest confidence reports possible, and we propose that the belief g at decision time is mapped onto this interval in a deterministic way. Thus, we are able to compute the probability density for responses times, performance and confidence in 2AFC tasks with time varying evidence. For static stimulation, we assume that subjects internally sample the presented stimulus, and thus perceive a time varying stream of evidence with an unknown internal variance rate. Thus, we treat static and dynamic stimulation tasks in the same way.

## 2 Results

### 2.1 Model agreement with data

Our goals are to accurately model behavioral data in tasks with very different structure, mainly in tasks with static or dynamic stimulation with the same underlying optimal model for decision making, and moreover also be able to account for confidence judgements.

We use the three 2AFC tasks from (Ais et al., 2016) to study this problem. These are an auditory, contrast and luminance discrimination tasks. Shortly, the auditory and luminance tasks have dynamic stimuli, while the contrast task has static stimuli. The duration of the stimulation is different between tasks, namely the luminance stimuli are presented for 1 second and subjects are forced to decide in that interval, whilst the contrast and auditory stimuli last for 300ms but subjects can respond at any time. Furthermore, in the auditory task the pitch of two tones must be compared, thus there is a memory retention period of 500ms between the presentation of both tones (each lasts 300ms). However, we test whether the same model is able to explain the behavioral data in all of them.

Our modeling approach is to specify all the tasks in terms of the minimum amount of elements needed by an optimal decision agent to be able to decide. For instance, if the evidence samples in favor of each alternative have a gaussian distribution, then the net evidence (the difference between the evidence of the competing alternatives) is also normally distributed. Thus, it is optimal to only encode the net evidence’s distribution and not encode two distributions separately for each alternative. This approach also implies that the only effect that the memory retention period has on the decision model is to add a fixed delay in between trials. Furthermore, in tasks with limited stimuli duration (auditory and contrast) nothing must be integrated after the stimuli vanishes, and the choice is forced based on the observed samples.

Our key assumption is that the model considers that the samples of net evidence which are observed come from a gaussian distribution with known variance rate. In the dynamic stimulation tasks, said variance rate encompasses the external variance from which the actual sensory samples are generate, and an internal variance rate that is hidden to the experimenter. In the static stimulation tasks, the stimuli have no external variance rate because they are fixed, but we assume that the sensory information in the static stimuli is sampled and generates a stream, corrupted by internal noise. We assume that this stream of samples are also taken from a gaussian with known variance rate. Thus, are model treats static and dynamic tasks on the same footing. However, we set the variance of the net evidence, *σ*^2^, as a free parameter that is fitted based on the behavioral data, thus we can separate the external and internal variability, at least for dynamic stimulation tasks.

Furthermore, Drugowitsch et al. (2012) assume, as is typically done in most decision making models, that additional delays in the response time arise because of processing that is unrelated to optimal decision making, for instance memory retrievals and comparisons after the stimuli vanish or decision transmission delays. We do the same here and assume that the combination of these delays results in a fixed non-decision time probability distribution, which for simplicity we assume to be a half gaussian bell with average *τ_c_* and dispersion *σ_c_*.

Furthermore, Drugowitsch et al. (2012) also assume that there is a probability, *p_po_*, that subjects become distracted and respond randomly in a trial. The remaining decision model parameter is the subject’s urgency, encoded as a cost function for the passage of time *c*(*t*) (for simplicity we assume it is constant).

Finally, our proposed extension is that confidence reports are simply a mapping, or read-out, from the bayesian belief that the decision was correct, *g*, to the response range. The functional form of this mapping is currently unknown and represents a major open problem in models of confidence, which is referred to as confidence calibration (Meyniel et al., 2015). The benefit of our modeling approach is that we can deterministically compute the model’s RT and confidence probability densities for any mapping. Thus we are able to compare the relative quality of different mappings and also assess whether they are in good agreement with the observed behavioral data, which is one of our main goals. In order to simplify, we compare two mapping functions:

1. 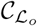, a combination of the widely used log probability ratio (logit of *g*) statistic (Kiani and Shadlen, 2009), with a sigmoid that rescales the reports to the interval [0, 1]. We refer to this mapping as the log-odds mapping.
2. 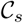, a linear rescaling of g, clipped to the [0, 1] interval. We refer to this mapping as the linear mapping.

We fit the extended model’s parameters (table 3) by maximizing the likelihood of the observed RT, performance and confidence for each subject, task and session independently (a total of 176 independent sets of parameters). We find that the model is able to fit the RT and accuracy distributions with great precision for all tasks (Fig. 1). This shows that the simplifying assumptions made to model the static and dynamic tasks on the same footing do not reduce Drugowitsch et al. (2012) model’s fitting power. This is important because in order to get good fits of the behavioral distributions in their original work, Drugowitsch et al. (2012) had to use a generic symmetric prior along with a time dependent cost function to get accurate fits, which added much more parameters to be fitted.

**Figure 1:**
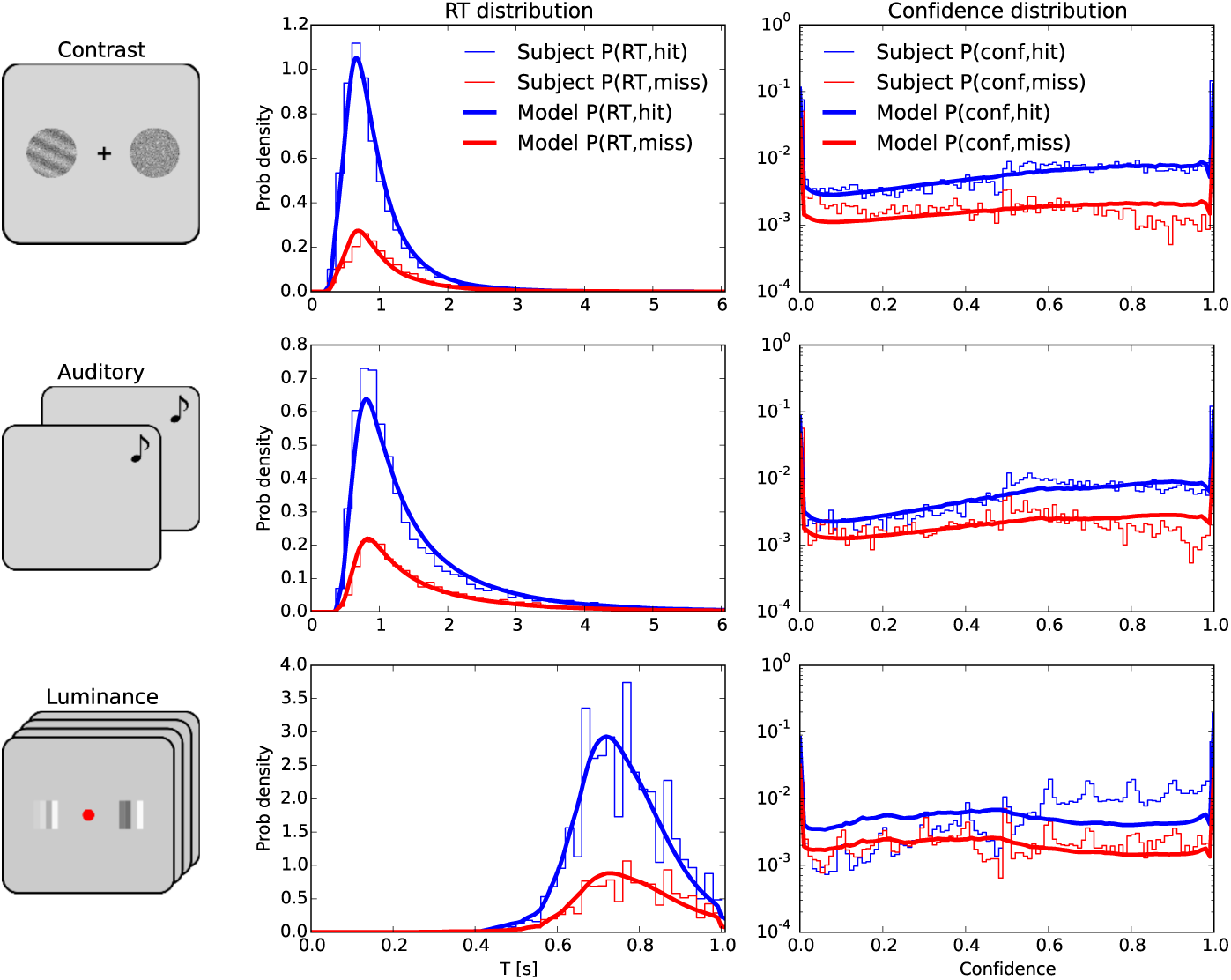
Model fit quality. We show the subject’s behavioral data: joint RT-accuracy distribution, confidence-accuracy distribution and binarized confidence-RT distributions, along with their corresponding fits for the 3 tasks. The subject’s confidence was binarized using a session mean split criteria.

Our proposed extension to model confidence, with the linear mapping, shows a good agreement between the response distributions for the auditory and contrast tasks (Fig. 1). However, the log-odds mapping does not yield a similar quality of the fits for both of these tasks (Fig. S.6). The proposed extension fails to fit the confidence distribution for the luminance comparison task, even though the data’s likelihood given the linear mapping is greater than given the log-odds mapping. We further asses this with a t-test to compare the difference between average confidence for each subject, session and experiment tuple, and its corresponding model fit. Grouping all the experiments together, the linear mapping shows no significant difference between the data and the theoretical prediction, while with the log-odds mapping, the model predicts significantly lower mean confidence than is observed (Table 1. For each experiment separately, the *T* statistics for the linear mapping show less significant differences than for the log-odds mapping.

**Table 1:** Average confidence difference t-test statistics for all experiments, Holm-Bonferroni corrected

| Experiment (mapping) | $n$ | $T$ | $p$ |
| --- | --- | --- | --- |
| All (linear) | 176 | -0.07 | 0.94 |
| All (log-odds) | 176 | -5.4 | $1.9 \cdot 10^{-6}$ |
| Contrast (linear) | 66 | 0.99 | 0.66 |
| Contrast (log-odds) | 66 | -2.3 | 0.08 |
| Auditory (linear) | 66 | 2.4 | 0.07 |
| Auditory (log-odds) | 66 | -2.8 | 0.03 |
| Luminance (linear) | 44 | -4 | 0.0015 |
| Luminance (log-odds) | 44 | -4.9 | $9 \cdot 10^{-5}$ |

We compare the relative quality of both mappings with the behavioral data’s maximum likelihood (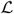). We find that for all experiments and session groups, the linear mapping has a better agreement than the log probability ratio mapping (Fig. 2). However, both mappings fail to explain the confidence resolution observed in the data as evidenced in the area under the receiver operating characteristic curves (AUC). Both mappings predict significantly lower AUC values (*p* < 10^−19^ for both mappings).

**Figure 2:**
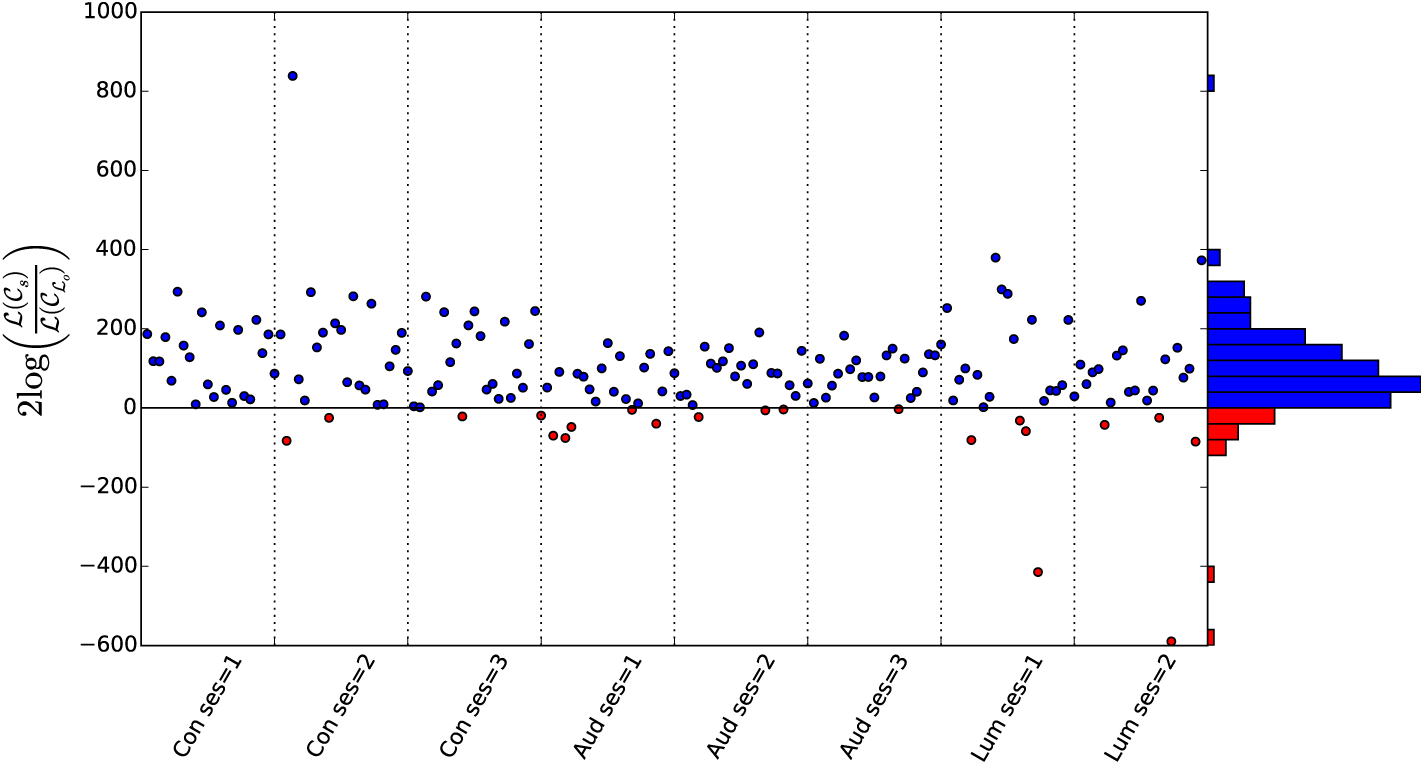
Comparison between mappings goodness of fit. Each dot corresponds to Wilks’ likelihood ratio statistic (Wilks, 1938) between both mappings for a single subject, session and experiment. Blue dots indicate that 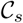 has a higher likelihood than 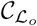. Red dots show the opposite case.

### 2.2 Parameter consistency

The model parameters determine the decision policy that will be followed. This means that by studying the fitted parameter values, we are able to asses whether the confidence mapping is the same for different tasks, and, more generally, if the confidence mapping is task idiosyncratic or if the readout policy is shared. Ais et al. (2016) did not observe significant learning across sessions, thus we hypothesize that policies across sessions of the same task should not vary greatly, but can significantly differ across tasks. This implies that parameter values of the same tasks should cluster together, and that clusters for different experiments may be separable.

To test this hypothesis, we take the fitted parameters for each subject, session and experiment. In order to make equally scaled parameters, we then normalize the values by the standard deviation of each parameter array, except for parameter *σ*^2^, where we use the standard deviation within each experiment for the normalization. We observe that parameter values for the same experiment, cluster together and that the different experiment clusters can be separated (Fig. 3.A and B). We also find that the clusters’ hierarchy, computed with Ward’s agglomerative clustering algorithm (Ward, 1963), shows distinct hierarchies for different parameter sets. The set of model parameters that determine the decision policy are the variance rate *σ*^2^, the cost *c*(*t*) and the phase-out probability *p_po_*. These parameters form a hierarchy where the auditory and luminance tasks policies are “closer” to each other than to the contrast task (Fig. 3.A). Whereas, for the confidence mapping parameters *C_H_* and a, the auditory and contrast tasks are closer than the luminance task (Fig. 3.B). In fact, Ais et al. (2016) observe a similar hierarchy for the confidence response distributions across tasks and sessions. They found that the auditory and contrast tasks confidence response distributions were more similar to each other than to the luminance task’s distribution.

**Figure 3:**
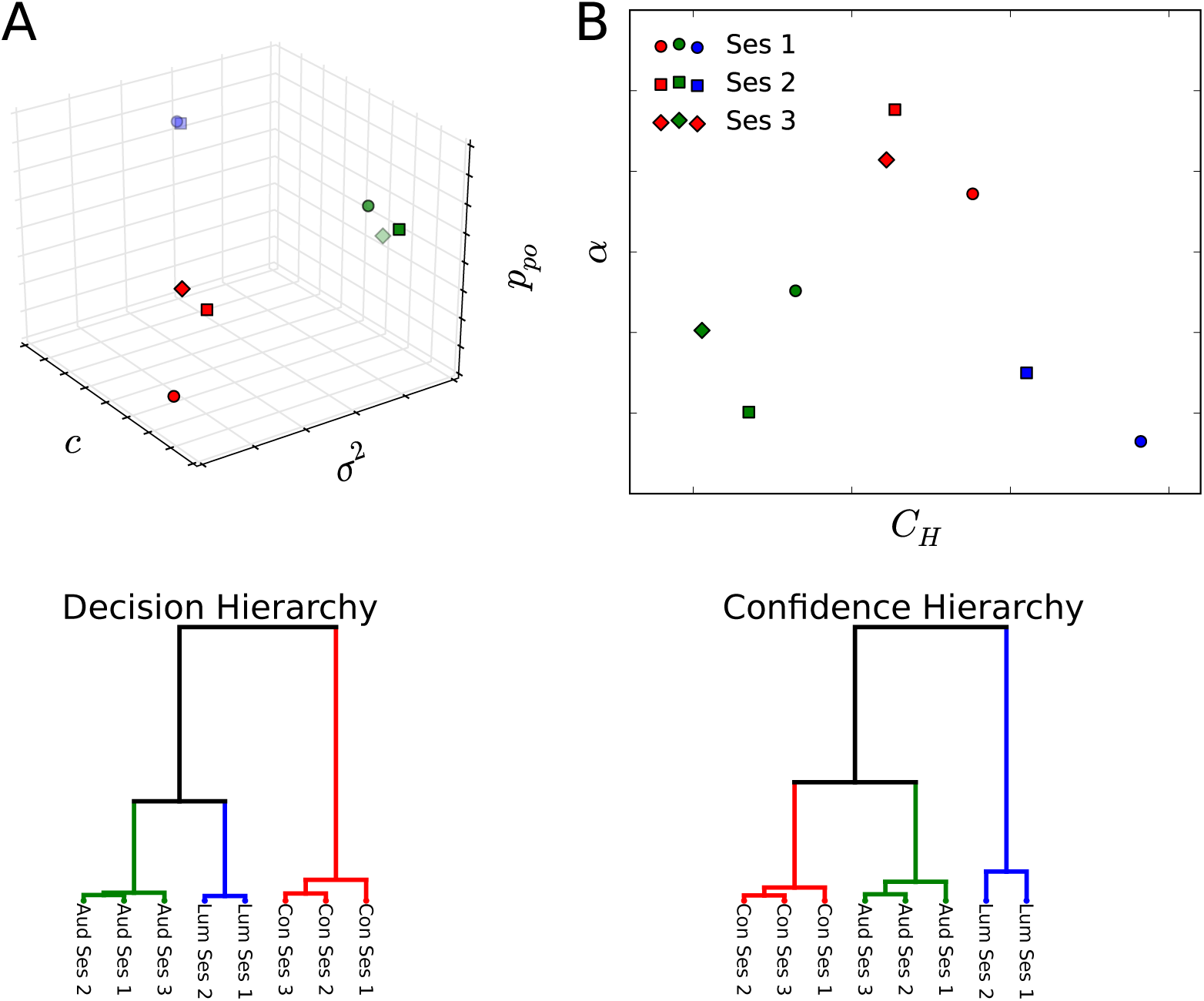
A shows a scatter plot of the relevant decision parameters averaged across all subjects for separate tasks and sessions. B shows is the same as A but for the relevant confidence parameters. The color of the markers label their corresponding task (blue=Luminance, red=Contrast, green=Auditory). Below each scatter plot, the hierarchy of clusters for the parameters (decision relevant A and confidence for B) is shown.

This hierarchy appears to be robust because most of the decision policy parameters are not significantly correlated to the confidence mapping parameters (*p* > 0.05 after holm-bonferroni correction). The significant correlations are between *σ*^2^ and *α* (corr= 0.25 *p* = 0.017), and *p_po_* and *C_H_* (corr= 0.31 *p* = 5.10^−4^). The former occurs because for large *σ*^2^ values, the belief bounds have a smaller slope (Sup. Fig. S.5), thus the mapping slope must be higher to be able to account for responses that span the entire response range [0, 1]. The latter correlation occurs because increased *p_po_* (more random response trials) is accompanied by lower confidence responses, thus the belief threshold to respond high confidence, *C_H_*, must be higher, which in turn explains the correlation. Thus, the only relevant correlations arise because of the model’s construction, leaving the rest of the parameters almost independent from each other.

## 3 Discussion

Our main goal is to test if confidence can be accurately be explained as a mapping of the bayesian posterior probability of having made the correct choice, as is proposed in many recent works (Meyniel et al., 2015, Pouget et al., 2016). We also aim to do this for tasks with different modalities, static and dynamic stimulation, to show that a general mapping principle can underlie several tasks response policies, but potentially with different mapping coefficients.

In order to do this, we study Ais et al. (2016) data for the three 2AFC tasks. The contrast task was completely static, while the other two were dynamic. The luminance task had a hard RT barrier, while the other tasks allowed subjects respond at any time, even after the stimuli vanished. Our modeling approach builds on Drugowitsch et al. (2012) decision making model and is able to accurately reproduce subjects RT distributions and performance for all three tasks. This is a crucial result because of the many simplifying assumptions that were made for fitting the model parameters. In their original paper, Drugowitsch et al. (2012) were able to fit behavioral distributions by allowing *c*(*t*) to vary with time and by using discrete symmetric priors on *μ*, instead of constant cost and conjugate gaussian priors. Furthermore, they only tested the model for a task with a single evidence source. On the other hand, we used three tasks with different modalities, two sources of evidence and different sensory exposure times. We found that the simplest form of inference (using gaussian conjugate priors), the simplest *c*(*t*) functional form, and making several assumptions on optimality (subjects compute net evidence, all delays of the decision beyond the stimulation duration comes from a fixed non-decision time) are sufficient to accurately explain the RT and performance for all tasks, static and dynamic stimulation alike.

Our main contribution is that we propose a way in which to compute the model’s predicted confidence responses based solely on the bayesian posterior belief. We compare two mappings from bayesian belief to confidence: the logit of the belief, mostly referred to as the log-odds, and a linear mapping from belief to confidence. We find that the linear mapping has better agreement for 87.5% of the subject, session and task tuples studied. Furthermore, we find that our proposal has a very good agreement for the auditory and contrast tasks, while the luminance task’s confidence response distribution is not well explained. We consider that this must be caused by one of simplifying assumptions not being true. One clear source of errors is that subjects rely on heuristic policies to report confidence (Maniscalco et al., 2016, Zylberberg et al., 2012), in combination to optimally inferred estimates (Paz et al., 2015). These imperfect decision policies produce performance drops that the model is able to explain at the expense of changing some parameters as the variance rate *σ*^2^. This can lead to an altered belief and bound form that makes it impossible to account for the confidence response distribution, even though the RT and performance distributions match. Furthermore, it has been recently found that subjects sometimes use measures of the information carried by the stimuli in order to compute their confidence, instead of just relying on the inferred probability of having made a correct choice (Ahumada et al., 2017). This strategy would also lead to confidence responses that could not be efficiently modeled using just the mapping of posterior belief.

On the other hand, the model parameters encode the decision policy robustly. That is, in the absence of observable learning during the experiments, the parameters do not vary broadly across sessions of the same task and cluster together. Furthermore, for the data studied here, the parameters formed distinct hierarchies of clusters where sessions of the same tasks group together first and then inter-task connections were made. A surprising observation is that the parameter cluster hierarchy is different for different parameter classes. In other words, the parameters that determine the decision policy (*c*(*t*), *σ*^2^ and *p_po_*) have a different hierarchy than the confidence mapping parameters (*C_H_* and *α*). Furthermore, the confidence mapping parameters cluster in a hierarchy that is equal to the one observed by Ais et al. (2016) when they studied the confidence response distributions similarities. This seems to indicate that the model parameters can represent hidden decision policy features that are differently shaped by task structure.

If we consider the decision parameters’ hierarchy, the dynamic tasks are more closely related than the static task, while in the confidence mapping parameters’ hierarchy, the short presentation tasks are more closely related than the Luminance task. This appears to indicate that the decision policy is strongly altered by task modality (dynamic or static), while the confidence mapping is strongly altered by the stimulation duration. However, it is not possible to make a conclusive statement about these observations only based on the hierarchies of fitted parameters, since these could be affected by many artificial aspects such as the scaling method, the clustering algorithm used, and even by inter-parameter correlations (although we did not observe significant correlations for most of the decision-confidence parameter pairs). The fact that the hierarchies are different can however be taken as evidence for the dissociated nature of perceptual accuracy and confidence reports, as was already reported in several studies (Cortese et al., 2016, Graziano and Sigman, 2009, Grimaldi et al., 2015).

Our study contributes to the discussion of whether the same brain circuitry involved in decision making is used for the encoding of confidence. The optimal bayesian decision model derived by Drugowitsch et al. (2012) has a one-to-one mapping between the integrated sensory evidence and the belief of having made the correct choice, as a function of elapsed time. We propose that confidence is then read-out from the bayesian belief, thus the circuitry for decision and confidence must be shared up until the read-out circuit. The read-out is implemented as a simple mapping from bayesian belief. We found that this mapping can be affected by task structure, which would lead to possible dissociations with actual decision performance.

## 4 Methods

### 4.1 Experimental data

We study the behavioral data measured by Ais et al. (2016) in three 2AFC tasks: an auditory discrimination task, a contrast discrimination task and a luminance discrimination task. In the auditory task, subjects had to listen to two successive pure tones corrupted by gaussian noise and had to indicate if the first or second one had a higher pitch. Both tones lasted 300ms and they were separated by a 500ms interval. In the contrast task, subjects had to maintain their fixation on a central cue, while two circular stimuli appeared to the sides for 300ms. One of them held a Gabor grating of random orientation superimposed with noise while the other only had noise, and the subjects had to identify which target contained the grating. In the luminance task, subjects had to fixate on a central cue while two patches of bars with different brightnesses flickered to the sides for 1 second. The bars brightness changed each 50ms around an average luminance, and the subjects had to identify which patch had higher mean brightness.

In all the tasks, subjects also reported their confidence in a continuous scale. The difficulty of the tasks was adapted with a Quest staircase (Watson and Pelli, 1983) to stay in the range of 75%. The auditory task allowed the subjects to decide after the stimuli had disappeared, and they could take all the time they needed to decide. The contrast task allowed the subjects to respond at any time, even before the stimuli had vanished. However, the luminance task forced the subjects to decide in the first second of stimulation.

The contrast task had completely static stimuli while the luminance task had large imposed dynamic noise. The auditory discrimination task imposed gaussian spectral noise with an amplitude that was 80% smaller than the pure tone’s amplitude.

The data we model are the subject’s decision times, performance and confidence, across different tasks and sessions. We use the data from 22 subjects who performed all 3 sessions of the auditory and contrast tasks, and the 2 sessions of the luminance task. We also use information of the observed task difficulties of each subject and session, for the model’s construction.

### 4.2 Decision model

Drugowitsch et al. (2012) propose a model for 2AFC tasks, where the subjects must select between two competing hypotheses (*H*_1_ and *H*_2_) based on a continuous stream of net evidence samples {*δx*_0_, *δx*_2_, … *δx_T_*}, where *δx*_t_ is the net evidence observed in the interval [*t*,*t* + *δt*). The samples of evidence are assumed to be generated from a normal distribution with unknown mean but known variance, *N*(*μ, σ*^2^). The goal of the model is to decide which of the underlying hypotheses generated the net evidence stream. The two competing alternatives are if *μ* >= 0 (*H*_1_) or *μ* < 0 (*H*_2_). Thus, the model must compute the belief it has that, for instance, *H*_1_ generated the observed evidence stream. Using bayes rule and assuming a conjugate prior, this can be written as:

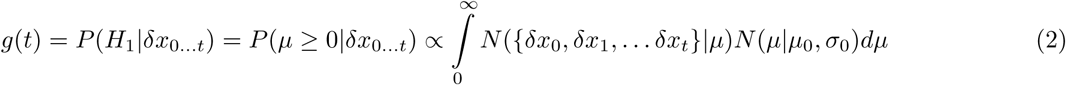
where *N* is the normal probability density, and *μ*_0_ and *σ*_0_ are the prior distribution’s hyperparameters. The belief that the alternative hypothesis is correct is simply 1 − *g*(*t*), because both hypotheses are mutually exclusive.

Being able to compute its belief in favor of each alternative, the model must establish a decision criterion that tells it when its belief is strong enough to commit to a decision. In order to do this, Drugowitsch et al. (2012) propose that the model’s goal is to maximize its reward rate: 
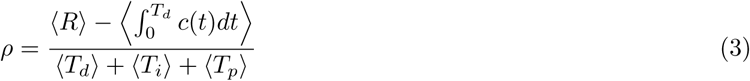
where *R* is the reward received after a decision, *c*(*t*) is the cost incurred at time *t* during the decision process, *T_d_* is the decision time, *T_i_* is the inter-trial interval, *T_p_* is extra penalty time added after an error trial, and the averages ⟨·⟩ are taken over all trials. Drugowitsch et al. (2012) show that in order to maximize the reward rate, the model must compute the average adjusted value for holding belief *g* at time *t*, 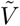 (*g, t*) (Mahadevan, 1996). This is done by solving the following Bellman equation: 
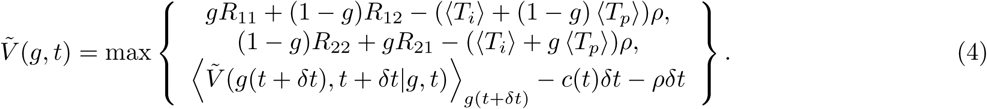
where *R_i,j_* is the reward obtained for selecting alternative *j* when *i* was correct. The first two terms compared in the max operation represent the average adjusted value associated to deciding immediately in favor of *H*_1_ and *H*_2_ respectively, while the last term is the value of waiting some more before deciding. Because the alternatives are equally likely in the tasks we analyze, and no penalty time is added, we simplify the above expression by assuming *R*_11_ = *R*_22_ = 1 and *R*_12_ = *R*_21_ = 0 and ⟨*T_p_*⟩ = 0. The Bellman equation is then solved by discretizing the belief interval in *n* bins, setting a maximum time *T* where it is not possible to delay the decision and propagating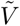 (*g, t*) from 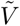 (*g,T*) backwards. For the detailed computation of 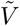 (*g,t*)refer to supplementary information sec. 1. From 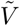 (*g,t*) it is possible to compute the bounds in belief space *θ*_1_(*t*) and *θ*_2_(*t*) where it becomes more valuable to decide immediately than to decide later (Fig. S.1.B and C). These bounds depend on the cost, *c*(*t*), variance *σ*^2^, prior hyperparameters and, most importantly, on time *t*. As time passes, the value of delaying the response falls because the expected future value 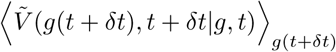 drops asymptotically to 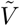 (*g, t*) with time (Fig. S.4). The cost function is able to modify the shape of the drop, but by no means is it imperative that *c*(*t*) ≠ 0 for the bounds to drop to zero.

Thanks to the bijective relation between belief *g* and accumulated net evidence *x*(*t*) = ∫ *δx_t_* (eq. 2), bounds *θ*_*i*_(*t*) can be transformed to *x*(*t*) space, *θ*_*xi*_(*t*). This enables us to compute the probability density of the evidence accumulating to the bounds as a function of time, *p*_*i*_(*t*) (FPT for first passage time, refer to supplementary information sec. 1 for details on the computation). By convoluting the FPT probability density with a non-decision time density, which incorporates all other delays not associated to the evidence integration process that occur between starting to perceive the stimuli and reporting the decision, we can obtain the model’s predicted RT probability density. We assume a simple half gaussian shape for the non-decision time density, *w*(*t*): 
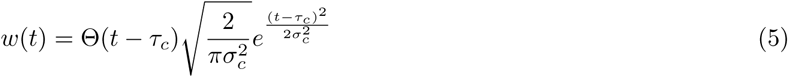
where *τ*_*c*_ is the average non-decision time, *σ*_*c*_ is the dispersion and Θ is the Heaviside step function.

We also assume that subjects have a probability *p_po_* of being distracted and making a random report in a given trial. We assume that the RT probability density for these “phased-out” trials is a uniform distribution between RT_min_ and RT_max_, the minimum and maximum RTs in the session. Thus the model’s predicted RT is 
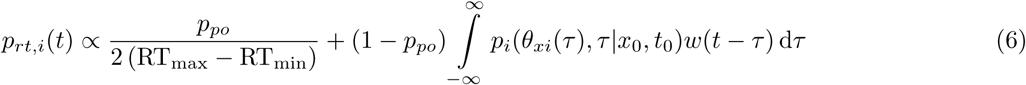
where *p_i_*(*θ*_*xi*_(*τ*),*τ*|*x*_0_,*t*_0_) is the FPT density of having first reached threshold *θ*_*xi*_(*τ*) at time *τ* and having started the integration at time *t*_0_ and evidence *x*_0_.

### 4.3 Confidence extension

The model determines the decision once the belief, *g* or 1 − *g*, that one of the competing alternatives is correct reaches a bound. This belief is nothing more than the probability that the decision taken is correct. We assume that subject’s reported confidence is computed as a mapping from belief *g* or 1 – *g*, to the response interval 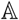, which we assume to be the closed interval [0, 1], where 0 and 1 represent the lowest and highest available confidence responses respectively. We explore two possible mappings. The first is the widely used logit of the selected alternative’s belief, most commonly referred to as the log probability ratio: 
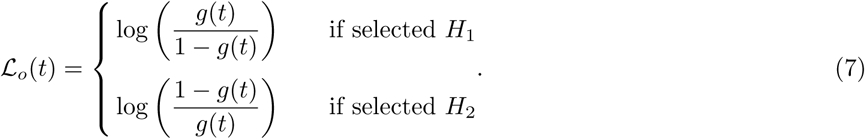

This mapping simply computes the logarithm of the ratio of the probability of having decided correctly over the probability that the alternative was correct. The variable 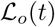 lies in the interval [0, + ∞) for the tasks that we study, where both alternatives are equally rewarded and, a-priori, equally likely. Thus it must then be remapped to the response interval [0, 1]. We assume this is done with a logistic function 
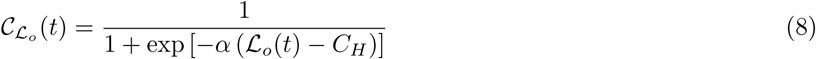
where *α* is the transition slope and *C_H_* is the high confidence threshold. The second mapping we test is a simple linear mapping 
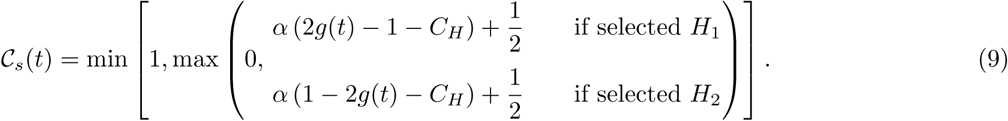

The operations 2*g*(*t*) − 1 and 1 − 2*g*(*t*) are performed in order to rescale the belief for each selection to the interval [0, 1] where 0 is low belief and 1 is high belief. Before doing this, for *H_1_* decisions *g*(*t*) lies in the interval [0.5, 1], where 0.5 and 1 correspond to low and high belief respectively, while, for *H*_2_ decisions, *g*(*t*) is in the interval [0,0.5] where 0 and 0.5 correspond to high and low belief respectively. The min and max operations are done to clip the value of *C*_*s*_(*t*) to the interval [0, 1]. The parameters *α* and *C_H_* have the same interpretation as in eq. 8 (Fig. 4.A). It is worth to note that when *α* → + ∞, both mappings produce binary confidence reports. This implies that in order to study datasets where subjects responded only binary confidence values, it is necessary to set *α* → + ∞ (Fig. 4.A).

**Figure 4:**
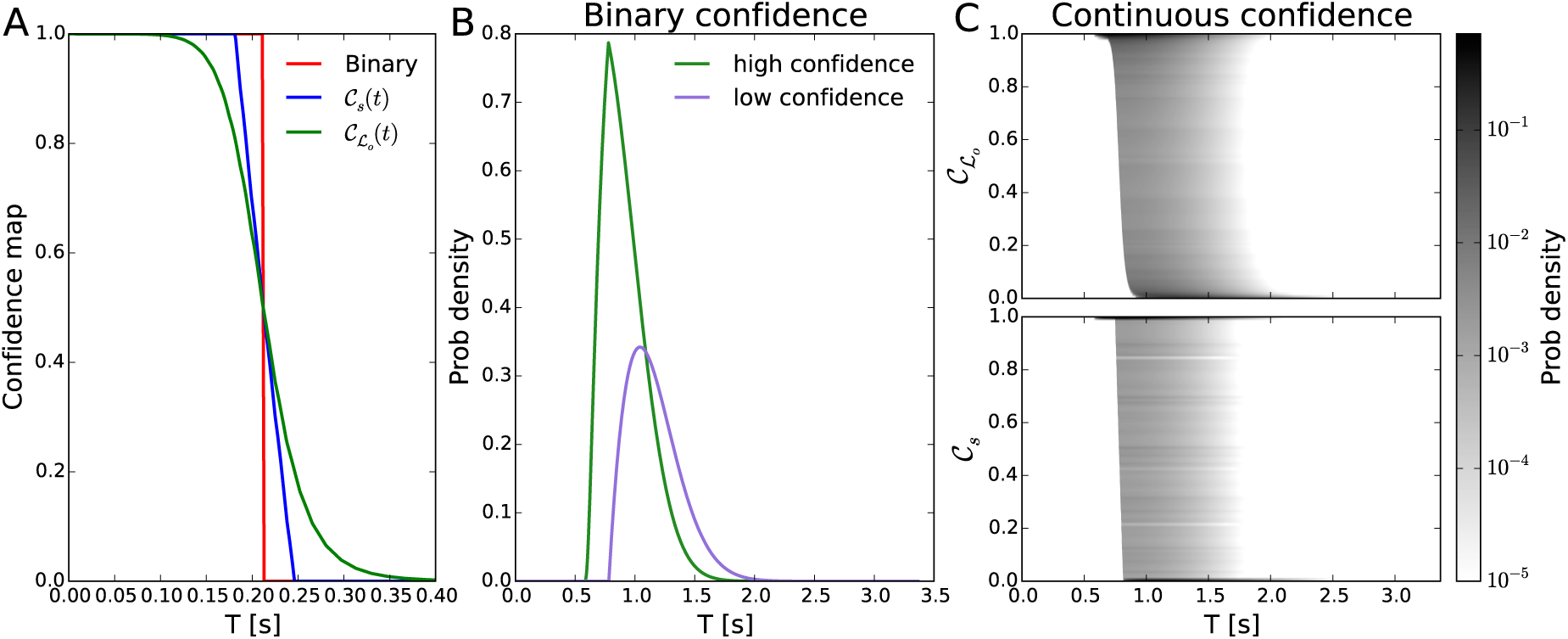
We show a sketch of the belief to confidence mapping. A. The resulting confidence response for both mappings calculated for the same belief data. The binary response curve (red) is computed setting *α* → ∞. B. shows the predicted high and low confidence response time distributions for the binary confidence response. C. shows the joint distribution of RT and confidence responses for both of the belief to confidence mappings.

Both mappings assume that only the belief *g* is used to compute confidence. This implies that the decision bounds in belief space *θ*_*i*_(*t*) determine confidence in a deterministic way. Thus, analogously to eq. 6, using the first passage time probability density *p*_*i*_(*t*) and the non-decision time distribution, we can compute the probability density of responding at time *t*, for alternative *i* with confidence co as: 
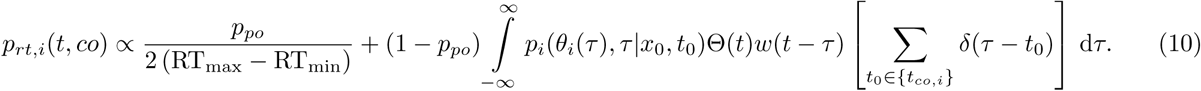
{*t_co,i_*} is the set of times where the confidence mapping returns the co value (*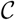*(*t*_*co,i*_) = *co*) for the decision i (Fig. 4.B and C).

### 4.4 Parameter fits

The decision model is controlled by several parameters most of which are completely determined by the task structure and some of which are fitted to account for the behavioral data. The former group is made up by all the parameters that are related to the reward rate eq. 3, the prior distribution hyperparameters and the model’s discretization parameters (*T, dt* and *n*). A list of these parameter values is shown in table 2. The only problem arises with the prior probability density. We assume that the net evidence has a gaussian likelihood. This is true for the luminance and auditory tasks, but for the contrast task, noise is uniform binary and evidence cannot be defined properly. Nevertheless, in order to fairly compare all tasks, we assume that net evidence is sampled from a gaussian distribution with an unknown mean but with known variance. This variance is left as a free parameter that is fitted for each subject, session and task independently. Having assumed that all tasks have gaussian likelihood, it is very convinient to choose a conjugate prior because the posterior of bayes rule is also gaussian. However, the prior of the net evidence is not gaussian because the task difficulty is controlled by a Quest staircase (Watson and Pelli, 1983) in order to maintain a running performance. This leads to a bimodal distribution of trial mean net evidence *μ* (Fig. 5). However, we assume that subjects have a prior that is the best fitting gaussian of the observed mean net evidences of the session. As the alternatives are equally likely, the prior’s mean is always 0. The prior variance is set to the session’s observed sample variance: 
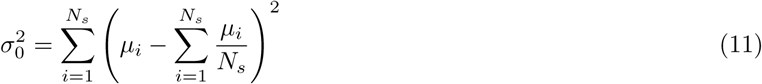
where the sums are performed over the observed *μs* for all trials in the session.

**Table 2:** Model parameters values fixed for each task, subject and session

| Parameter name | Luminance | Auditory | Contrast |
| --- | --- | --- | --- |
| $T$ | 1s | 0.3s | 0.3s |
| $\langle T_i \rangle$ | 1.5s | 2s | 1.5s |
| $dt$ | 8ms | 5ms | 5ms |
| $R_{11} = R_{22}$ | 1 | | |
| $R_{12} = R_{21}$ | 0 | | |
| $\langle T_p \rangle$ | 0s | | |
| $n$ | 101 | | |
| $\mu_0$ | 0 | | |
| $\sigma_0^2$ | From eq. 11 | | |

**Figure 5:**
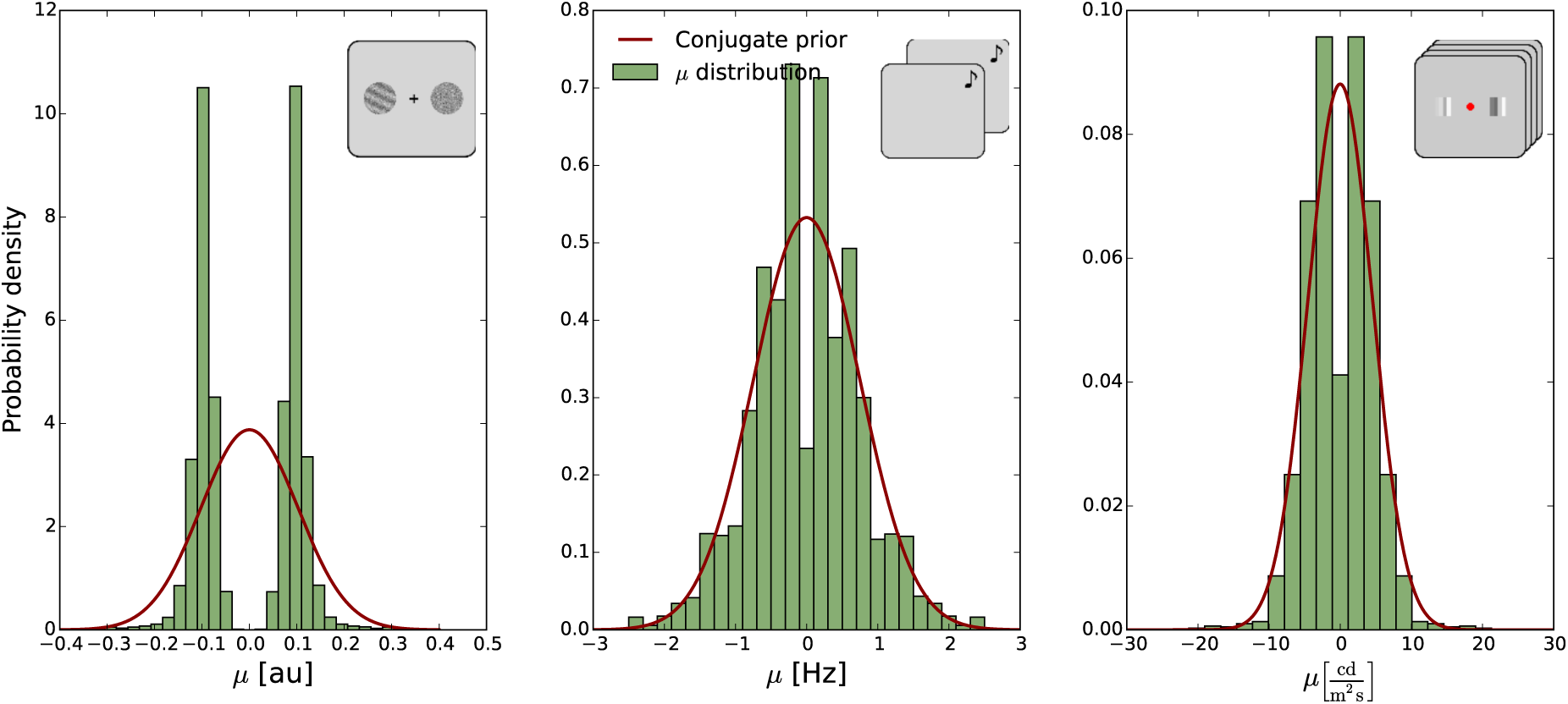
The prior distributions. We show the distribution of net evidence observed by the subjects (light green bars) and the corresponding gaussian fit. It is clear that all the distributions that the subjects viewed were bimodal, yet we assume that their internal prior distribution is a gaussian.

It is worth to note that *T* is set differently for each task because the optimal strategy differed depending on task structure. In the luminance task, subjects were forced to respond during the first second after stimulus onset. Thus, at time *t* = 1s, they were unable to delay their decision any longer. This is achieved by setting *T* = 1s. On the other hand, in the contrast and auditory tasks, subjects were able to decide at any time. However, the optimal decision strategy only takes into account the period of time the evidence is shown. Thus, we set *T* = 0.3s because it is the time the net stimulation lasts for both tasks. In the contrast task, the stimuli are presented in the screen for 0.3s and then disappear. In the auditory task, each tone is presented for 0.3s but the optimal decision strategy only depends on the difference between both evidence samples, thus it can be assumed that for the optimal decision maker, both stimuli arrive in parallel and they are presented for only 0.3s. The additional time taken for the decision report is assumed to be consumed by non-optimal decision related computations such as memory retrievals.

The free model parameters that remain are the cost of time (*c*(*t*)) which we assume to be constant, the generative model’s variance (*σ*^2^), the probability of “phase-out” trial (*p_po_*), the non-decision time mean and dispersion (*τ_c_* and *σ_c_*), and the confidence mapping threshold and slope (*C_H_* and α). All of these parameters were fitted to each subject, session and task independently. Using eq. (10) we are able to compute the likelihood of the observed RT, performance and confidence for the distribution of net evidence μ of each session. Thus, we determine the free parameters values as the maximum likelihood fit of the RT, performance and confidence. The fitted parameters are listed in table 3 along with their optimization procedure starting values and bounding regions.

**Table 3:** Model parameters that are fitted for each subject, task and session, along with their optimization procedure starting values and bounding regions

| Parameter name and description | Starting value | Searched interval |
| --- | --- | --- |
| $c(t)$, cost of passage of time | 0.02Hz | $[0, 0.4]$Hz |
| $\sigma^2$, generative variance | $\sigma_{st}^2$ | $[0.2\sigma_{st}^2, 1.8\sigma_{st}^2]$ |
| $p_{po}$, “phase-out” probability | 0.05 | $[0, 0.2]$ |
| $\tau_c$, non-decision time offset | 2.5% lowest RT | $[0, 1.5]$s |
| $\sigma_c$, non-decision time dispersion | $\sigma_{c_{st}}$ | $[0.001, 6]$s |
| $C_H$, high confidence threshold | $\mathcal{C} \left( \max_{rt} \{p_{rt,1}(t)\} \right)$ | $[0, M_C]$ |
| $\alpha$, confidence mapping slope | 17.2 | $[0, 100]$ |

The likelihood as a function of the parameters is very rugged and has several plateaus and local maxima that make it difficult to perform the fits. Furthermore, the likelihood depends on the computed decision bounds through parameters *c*(*t*) and *σ*^2^, thus we are unable to compute the gradient of the likelihood. Based on these limitations we use a covariance matrix adaptation (CMA-ES) genetic algorithm to fit the parameters (Hansen et al., 2009). However, this algorithm can also get caught up in local optimums if it is not supplied with a sufficiently good search region or starting point. We estimate *σ*^2^ as the value 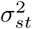 that better matches the subject’s performance for the observed drift rates *μ* based on a simplified logistic fit between performance and variance (see SI eq. S.33). We then bound *σ*^2^ to the interval 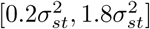(table 3). We set τ_*c*_ equal to the 2.5% lowest observed RT. The initial value of *σ_c_* is labeled as *σ_c/_st__* and computed based on the predicted probability distribution for the initial *σ*^2^ and *c*(*t*), *p_rt_*,i(t). We compute the RT variance of that distribution and subtract it to the subject’s RT variance. We set *σ_c/_st__* equal to 0.01s or to the squared root of the difference between the subject and model’s RT variance, if it is greater than 0.01s. Finally, the initial value of C_H_ is also computed from p_rt_,_1_(*t*). Given *p*_*rt*_,_1_(*t*) we compute the mode of the distribution, *t*_mode_ = max*_rt_p_rt_*,_1_(*t*), and set *C*_*H*_ = *C*(*t*_mode_). The search interval for C_H_ was set between 0 and the maximum value the used mapping could produce, 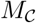. For 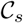, 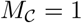 while for 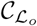, 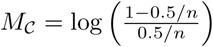. The rest of the starting points are set to values that are independent from the subject’s behavioral responses as can be seen in table 3.

### 4.5 Parameter clustering

The fitted parameter values determine the policy followed by the subjects. To test how much this policy varied across subjects, sessions and tasks we study their proximity as points in a multidimensional space, where each dimension corresponds to a different parameter. We assume that the distance in between points is euclidean, which allows us to use Ward’s agglomerative clustering algorithm (Ward, 1963). The only subtlety is that the scale, and moreover the units, of the parameters are very different. For instance, *c*(*t*) is measured in Hz, while *τ_c_* is measured in s. To make all scales comparable, we normalize the values of every parameter, with the exception of *σ*^2^, by their standard deviation across subjects, sessions and tasks. Parameter *σ*^2^ has to be treated differently because its scale and units are different for each task. In the auditory task, *σ*^2^’s units are Hz^3^, in the contrast task Hz and in the luminance task, cd^2^/m^4^Hz. Thus, we normalize the values of *σ*^2^ separately for each task dividing them by the standard deviation across subjects and sessions. This allows us to treat all the parameters on the same footing and consider that the normal euclidean distance can be used for the clustering. We use the python package scikit-learn (Pedregosa et al., 2011) to perform the clustering and compute the parameter hierarchy shown in Fig. 3. The hierarchy plots were performed using the ete3 package (Huerta-Cepas et al., 2016).

## 5 Acknowledgements

L. P. was supported by the Physics department and the Calculus Institute at the FCEyN, Universidad de Buenos Aires, and the Neuroscience Lab at the Universidad Torcuato di Tella. A.S. was supported by CONICET PIP 2014 1220130100384CO and UBACYT 2014 20020130200202BA. M.S was supported by CONICET, FONCyT Argentina Grant PICT-2013-1653 and the James S. McDonnell Foundation 21st Century Science Initiative in Understanding Human Cognition – Scholar Award. The Authors express that they have no conflict of interest with the published material.

